## Supplemental Information for "A nascent riboswitch helix orchestrates robust transcriptional regulation through signal integration"

**SUPPLEMENTARY FIGURES**

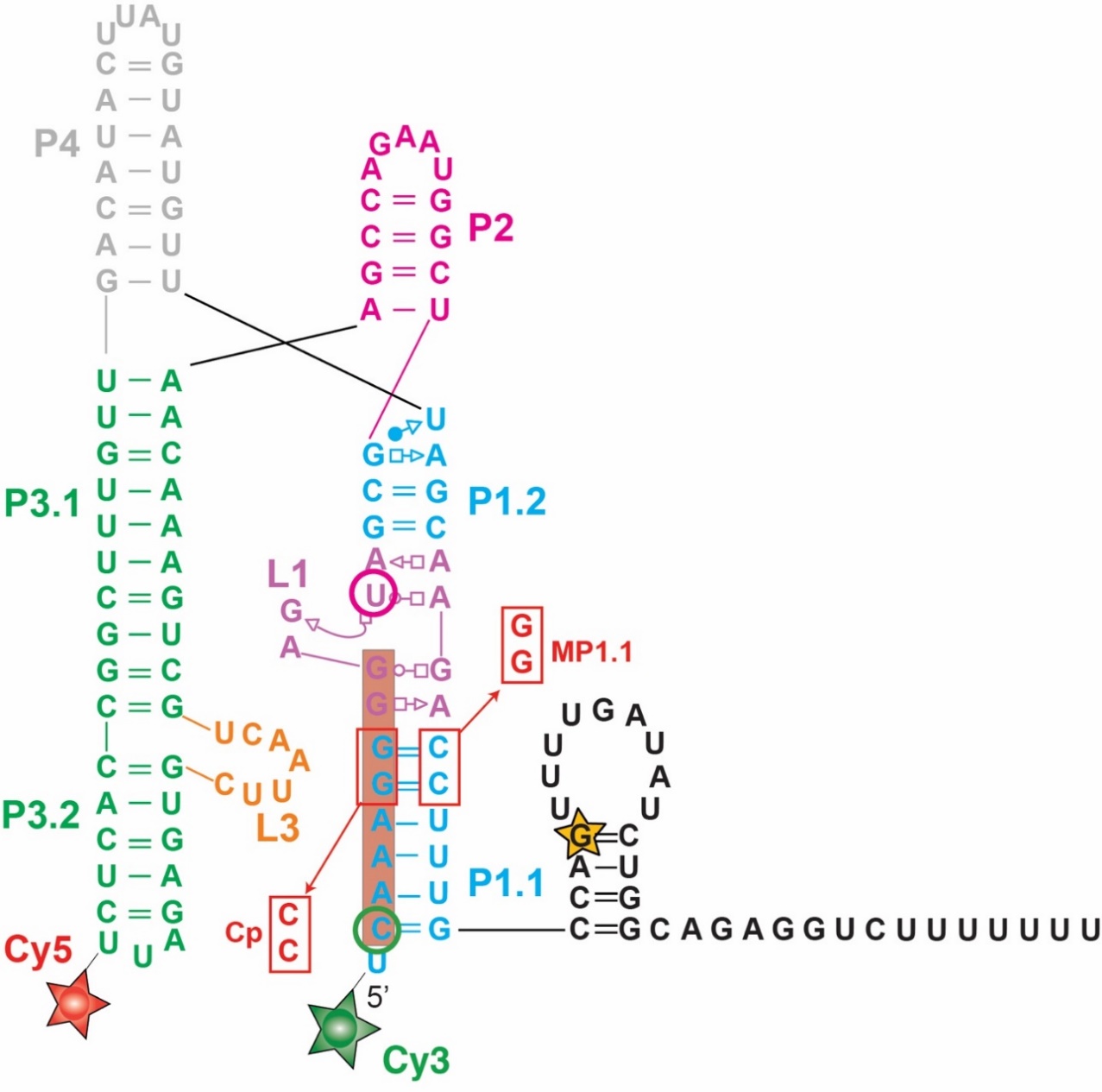

**Figure S1. Secondary structure of the *yybp* riboswitch**

The entire riboswitch sequence is shown with the P1 (cyan), P2 (pink), P3 (green), P4 (grey) stems, the L1 (purple), L3 (orange) loops and the terminator hairpin (black). RNAP pause at position G104 identified in this study is highlighted with a yellow star. The region targeted by the SiM-KARTS probe is shaded in dark orange. Positions of Cy3 and Cy5 fluorophores for smFRET study in nucleic scaffold assembly are indicated with green and red stars respectively. Positions of Cy3 and Cy5 fluorophores for smFRET study after stepwise transcription are indicated with green and magenta circles respectively Mutations destabilizing (MP1.1) and re-establishing (Cp) P1.1 are indicated in red.

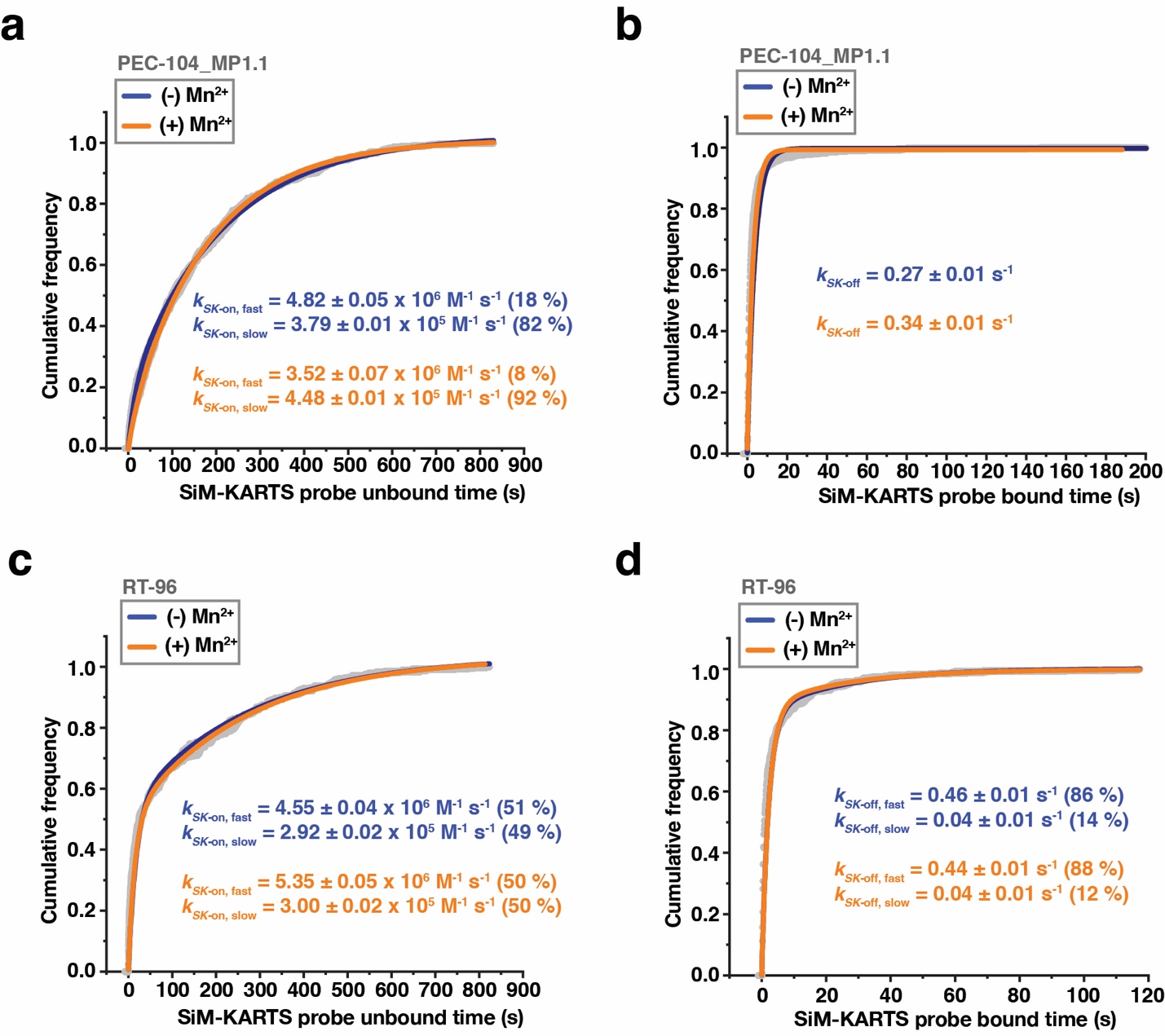

**Figure S2. SiM-KARTS probing of the P1.1 switching helix**

(a, b) Plots displaying the cumulative unbound (a) and bound (b) dwell times of the SiM-KARTS probe to the P1.1 in the context of PEC-104 – MP1.1. Binding (*k_SK_*_-on_) and dissociation (*k_SK_*_-off_) rate constants of the SiM-KARTS probe are indicated.

(c, d) Plots displaying the cumulative unbound (c) and bound (d) dwell times of the SiM-KARTS probe to the P1.1 in the context of RT-96. Binding (*k*_on_) and dissociation (*k*_off_) rate constants of the SiM-KARTS probe are indicated. Total number of molecules analyzed for each condition is as follow: PEC-104 MP1.1 (-) Mn^2+^ = 330; PEC-104 MP1.1 (+) Mn^2+^ = 292; RT-96 (-) Mn^2+^ = 118; RT-96 (+) Mn^2+^ = 182.

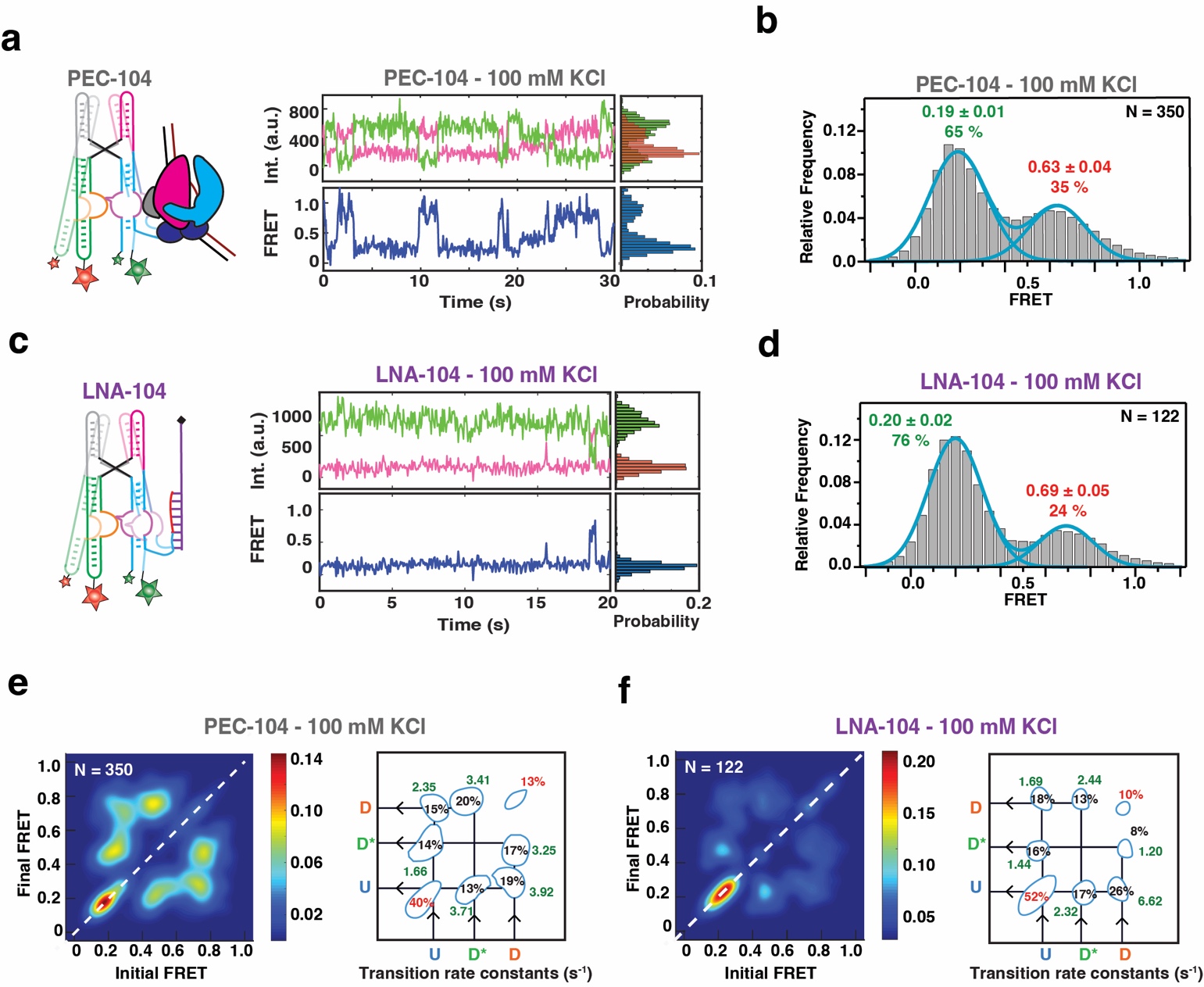

**Figure S3. smFRET-based docking analysis of the *yybp* riboswitch in the absence of divalent ions**

(a) Representative smFRET trace for PEC-104 in the absence of divalent ion (100 mM KCl). Green: Cy3, Pink: Cy5, Blue: FRET.

(b) Population FRET histogram showing the equilibrium distribution of two FRET states under the condition in panel a.

(c) Representative smFRET trace for LNA-104 in the absence of divalent ion (100 mM KCl). Green: Cy3, Pink: Cy5, Blue: FRET.

(d) Population FRET histogram showing the equilibrium distribution of two FRET states under the condition in panel c.

(e) TODP showing the static and dynamic traces as “on-diagonal” and “off-diagonal” heat map contours, respectively for PEC-104. The color code indicates the fraction of each population. Transition rate constants for each transition between the different FRET states are indicated on the right.

(f) TODP showing the static and dynamic traces as “on-diagonal” and “off-diagonal” heat map contours, respectively for LNA-104. The color code indicates the fraction of each population. Transition rate constants for each transition between the different FRET states are indicated on the right.

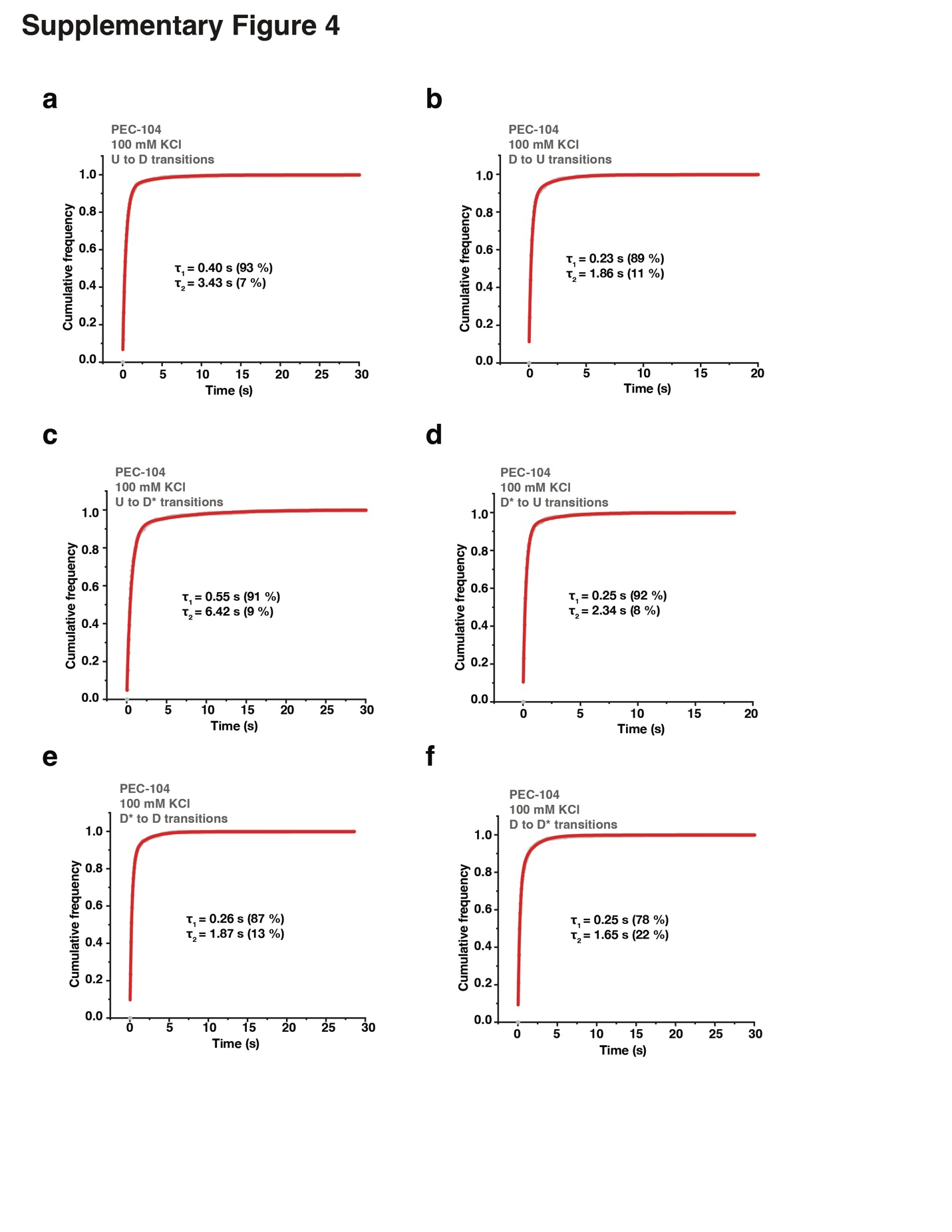

**Figure S4. Docking kinetics of PEC-104 in the absence of divalent ions**

(a-f) Cumulative dwell-time distributions of each transition between the three FRET states (U, D* and D) in the absence of divalent ions. The lifetimes and amplitudes of slow and fast components are also shown.

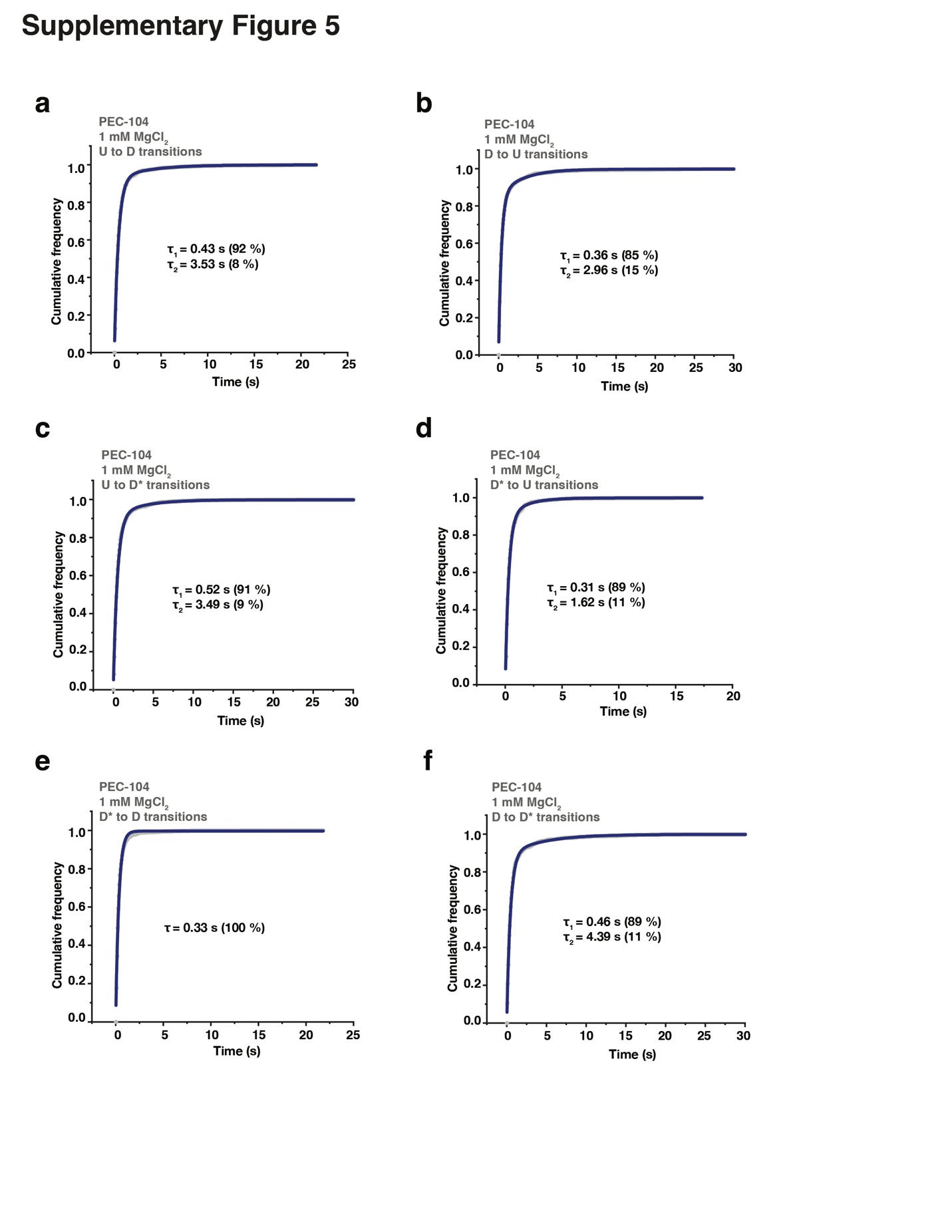

**Figure S5. Docking kinetics of PEC-104 in the presence of Mg^2+^ only**

(a-f) Cumulative dwell-time distributions of each transition between the three FRET states (U, D* and D) in the presence of 1 mM MgCl_2_. The lifetimes and amplitudes of slow and fast components are also shown.

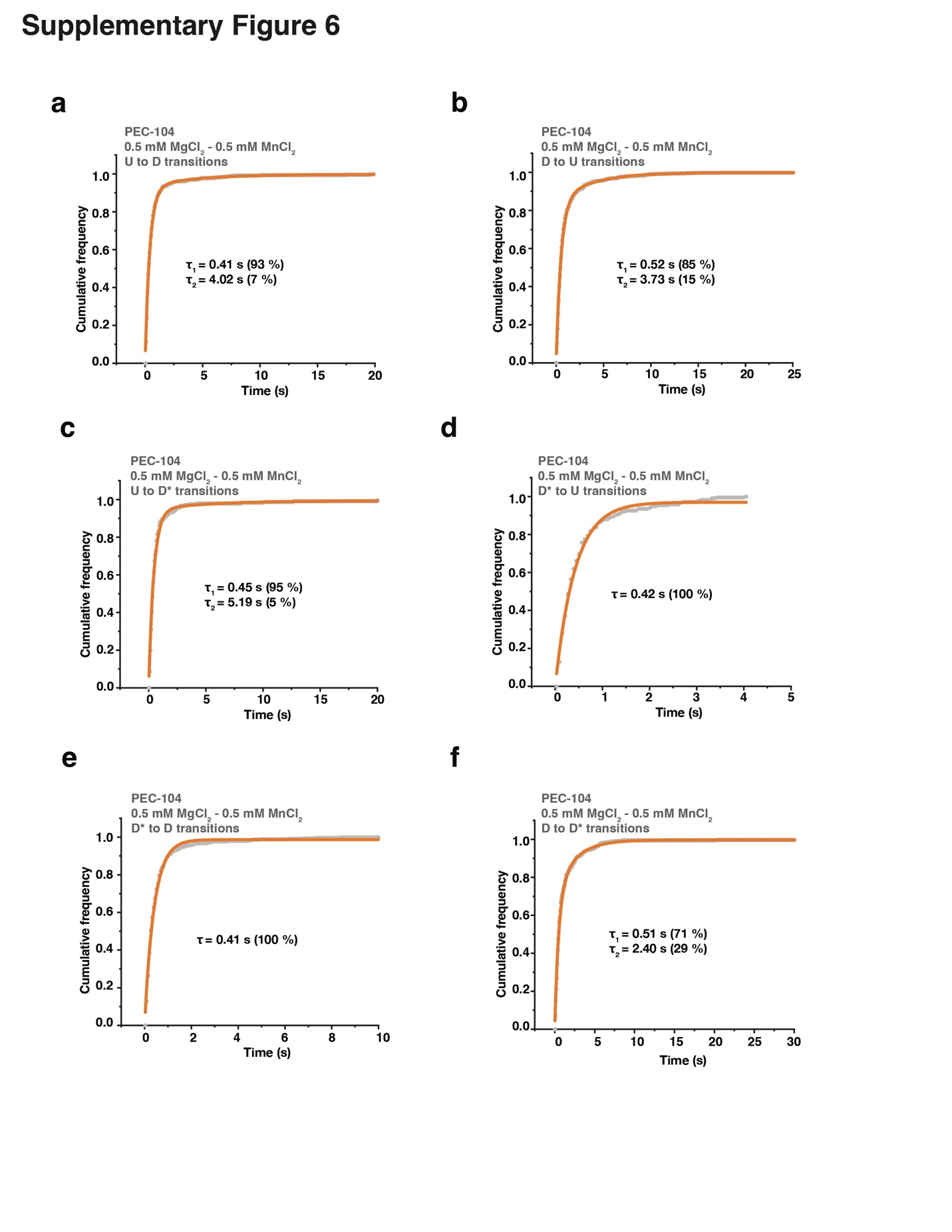

**Figure S6. Docking kinetics of PEC-104 in the presence of Mg^2+^ and Mn^2+^**

(a-f) Cumulative dwell-time distributions of each transition between the three FRET states (U, D* and D) in the presence of 0.5 mM MgCl_2_ and 0.5 mM MnCl_2_. The lifetimes and amplitudes of slow and fast components are also shown.

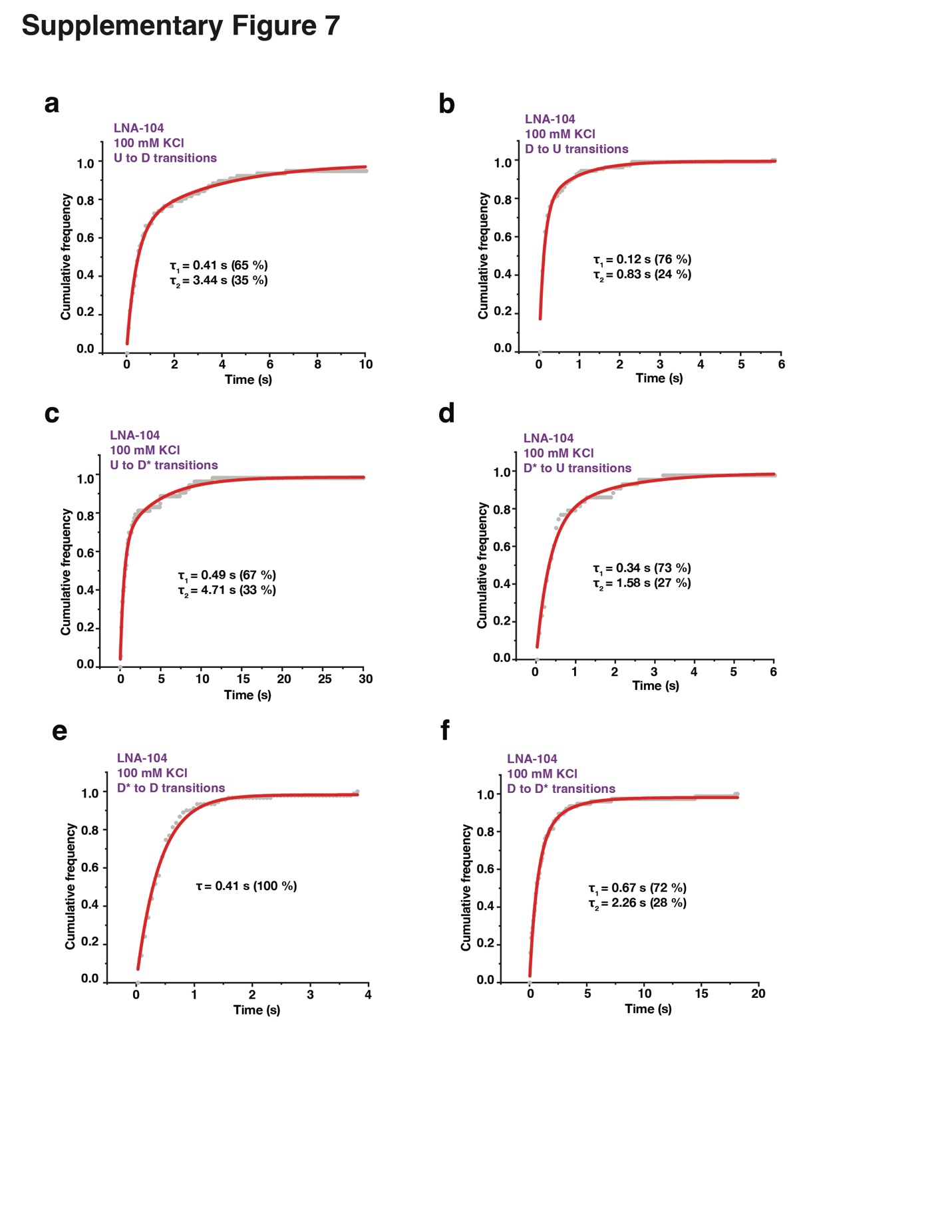

**Figure S7. Docking kinetics of LNA-104 in the absence of divalent ions**

(a-f) Cumulative dwell-time distributions of each transition between the three FRET states (U, D* and D) in the absence of divalent ions. The lifetimes and amplitudes of slow and fast components are also shown.

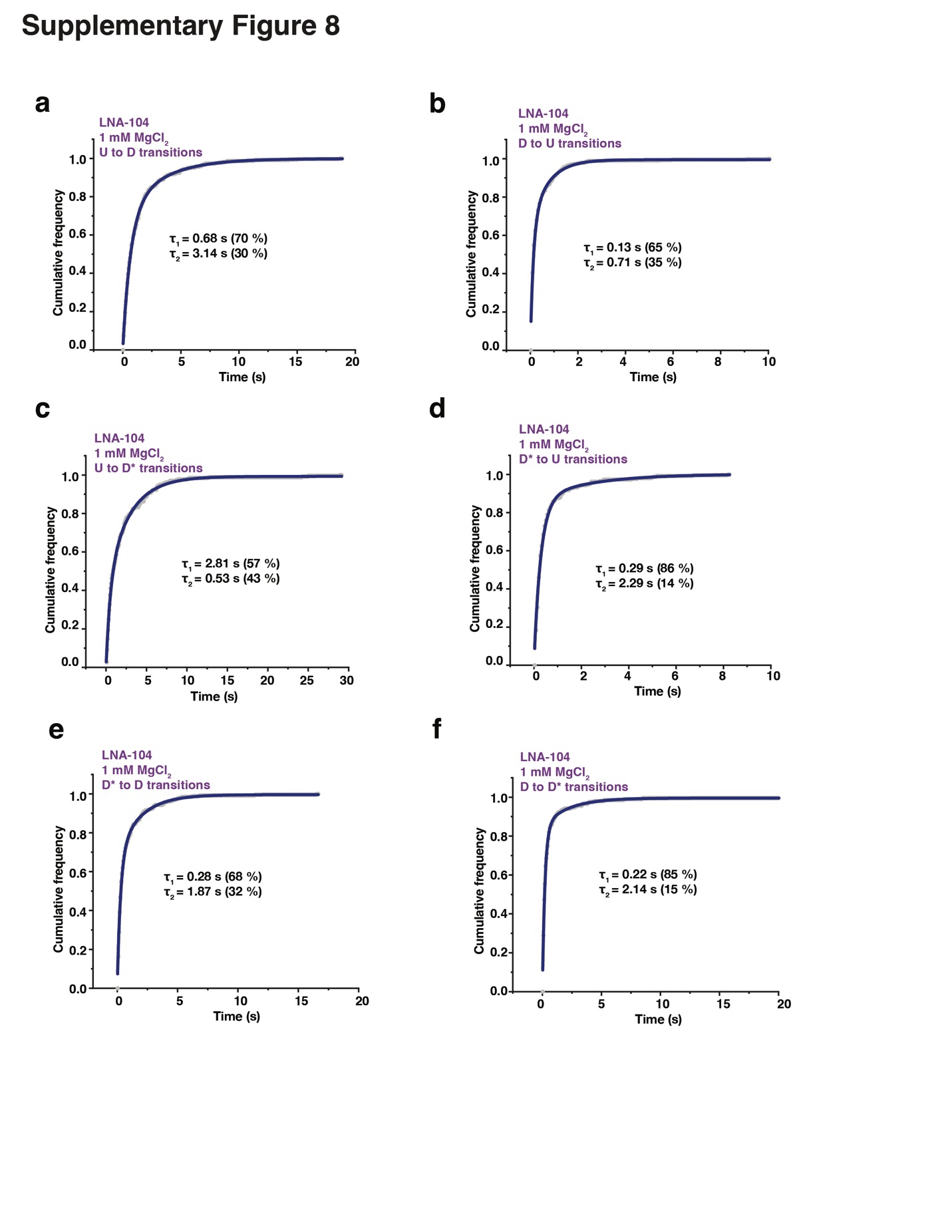

**Figure S8. Docking kinetics of LNA-104 in the presence of Mg^2+^ only**

(a-f) Cumulative dwell-time distributions of each transition between the three FRET states (U, D* and D) in the presence of 1 mM MgCl_2_. The lifetimes and amplitudes of slow and fast components are also shown.

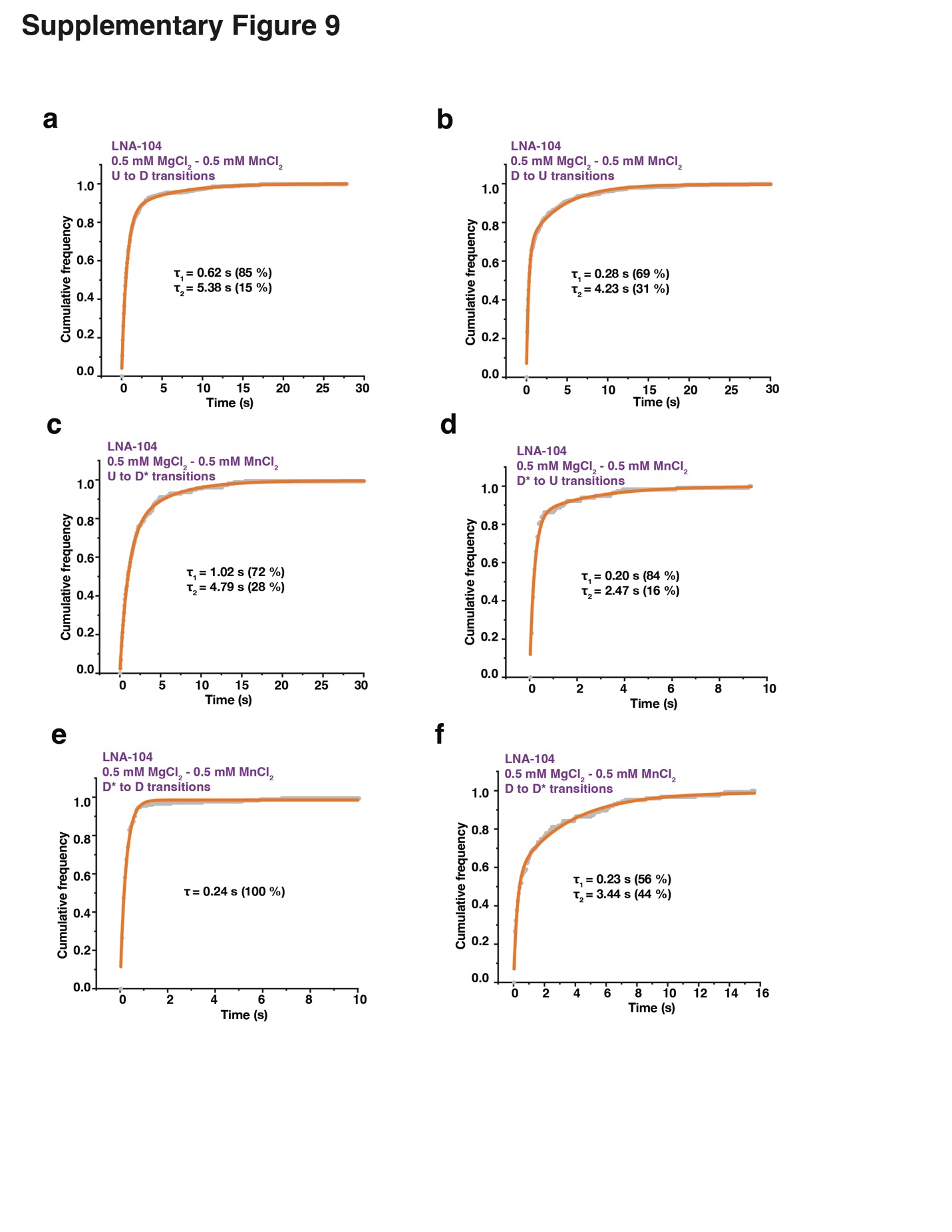

**Figure S9. Docking kinetics of LNA-104 in the presence of Mg^2+^ and Mn^2+^.**

(a-f) Cumulative dwell-time distributions of each transition between the three FRET states (U, D* and D) in the presence of 0.5 mM MgCl_2_ and 0.5 mM MnCl_2_. The lifetimes and amplitudes of slow and fast components are also shown.

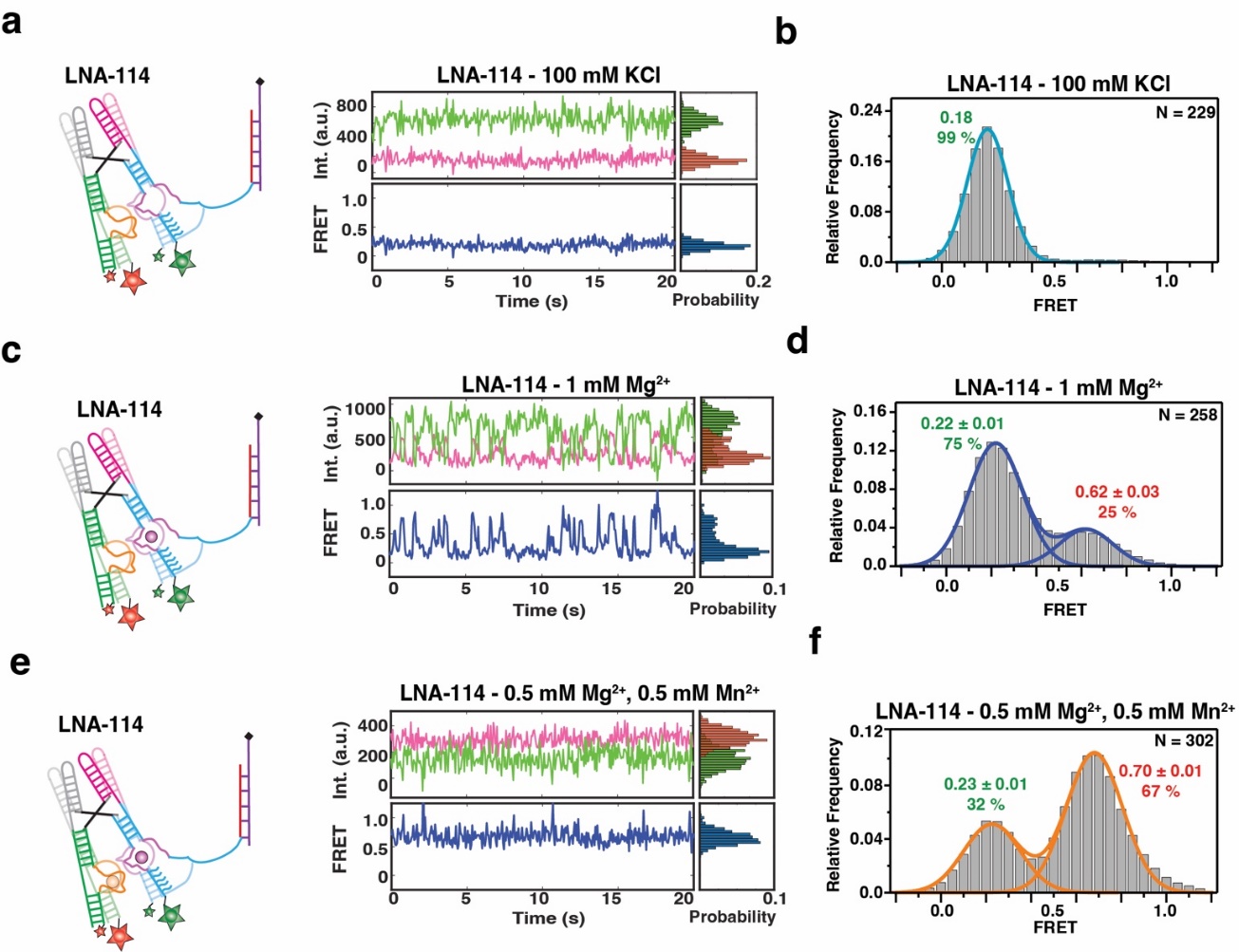

**Figure S10. smFRET-based docking analysis of the *yybp* riboswitch with a complete P1.1 helix**

(a) Representative smFRET trace for LNA-114 in the absence of divalent ion (100 mM KCl). Green: Cy3, Pink: Cy5, Blue: FRET.

(b) Population FRET histogram showing the equilibrium distribution of two FRET states under the condition in panel a.

(c) Representative smFRET trace for LNA-114 in the presence of Mg^2+^ only (1 mM Mg^2+^). Green: Cy3, Pink: Cy5, Blue: FRET.

(d) Population FRET histogram showing the equilibrium distribution of two FRET states under the condition in panel c.

(e) Representative smFRET trace for LNA-114 in the presence of Mg^2+^ and Mn^2+^ (0.5 mM Mg^2+^, 0.5 mM Mn^2+^). Green: Cy3, Pink: Cy5, Blue: FRET. (f) Population FRET histogram showing the equilibrium distribution of two FRET states under the condition in panel e.

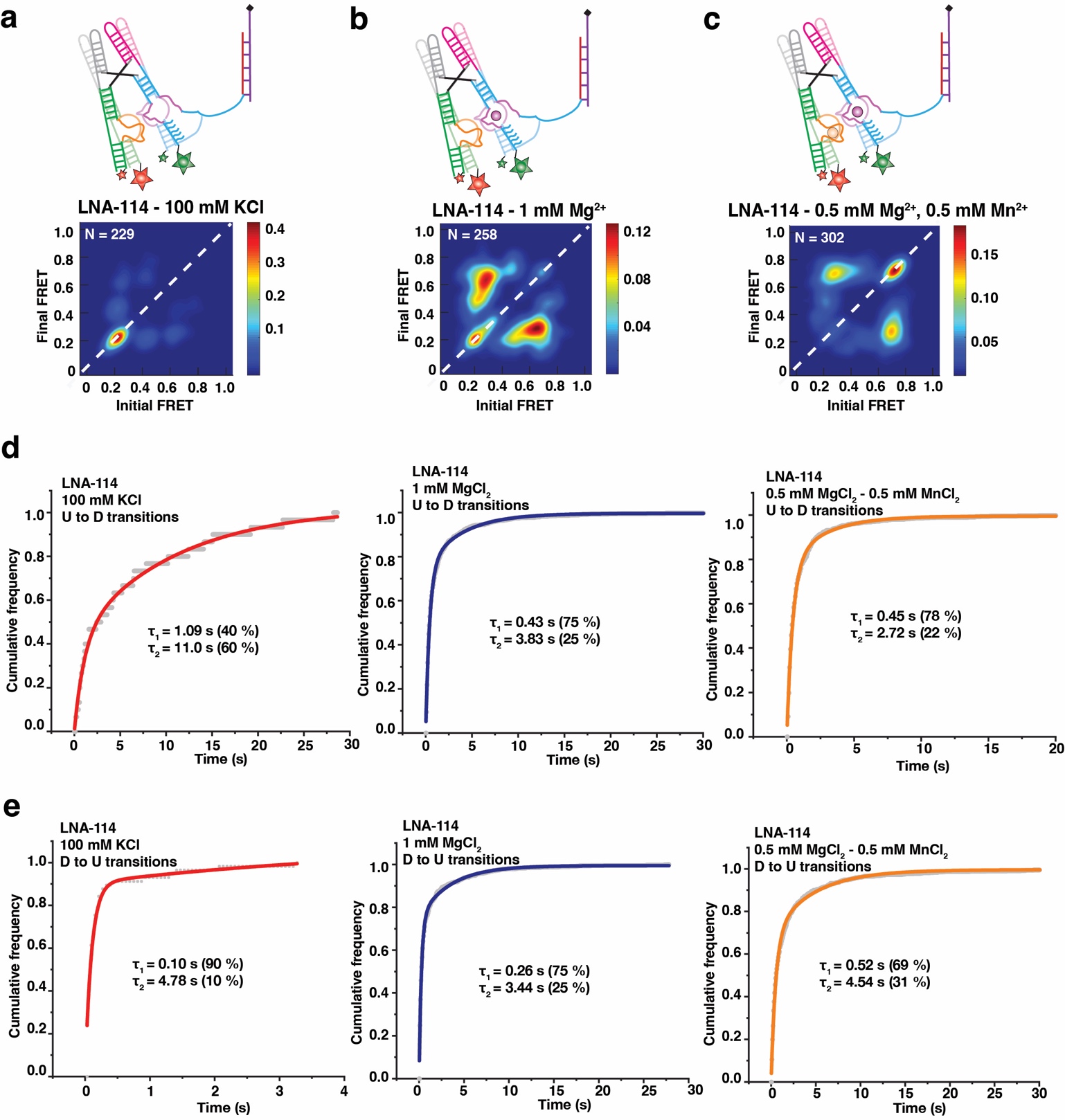

**Figure S11. Docking kinetics of LNA-114**

(a-c) TODP showing the static and dynamic traces as “on-diagonal” and “off-diagonal” heat map contours, respectively for LNA-114 under different conditions: a no divalent (100 mM KCl), b with Mg^2+^ only (1 mM Mg^2+^), c with both Mg^2+^ and Mn^2+^ (0.5 mM Mg^2+^, 0.5 mM Mn^2+^). The color code indicates the fraction of each population.

(d) Cumulative dwell-time distributions of the U to D state transitions in the absence of divalent (left), with Mg^2+^ only (middle) and with both Mg^2+^ and Mn^2+^ (right). The lifetimes and amplitudes of slow and fast components are also shown.

(e) Cumulative dwell-time distributions of the D to U state transitions in the absence of divalent (left), with Mg^2+^ only (middle) and with both Mg^2+^ and Mn^2+^ (right). The lifetimes and amplitudes of slow and fast components are also shown.

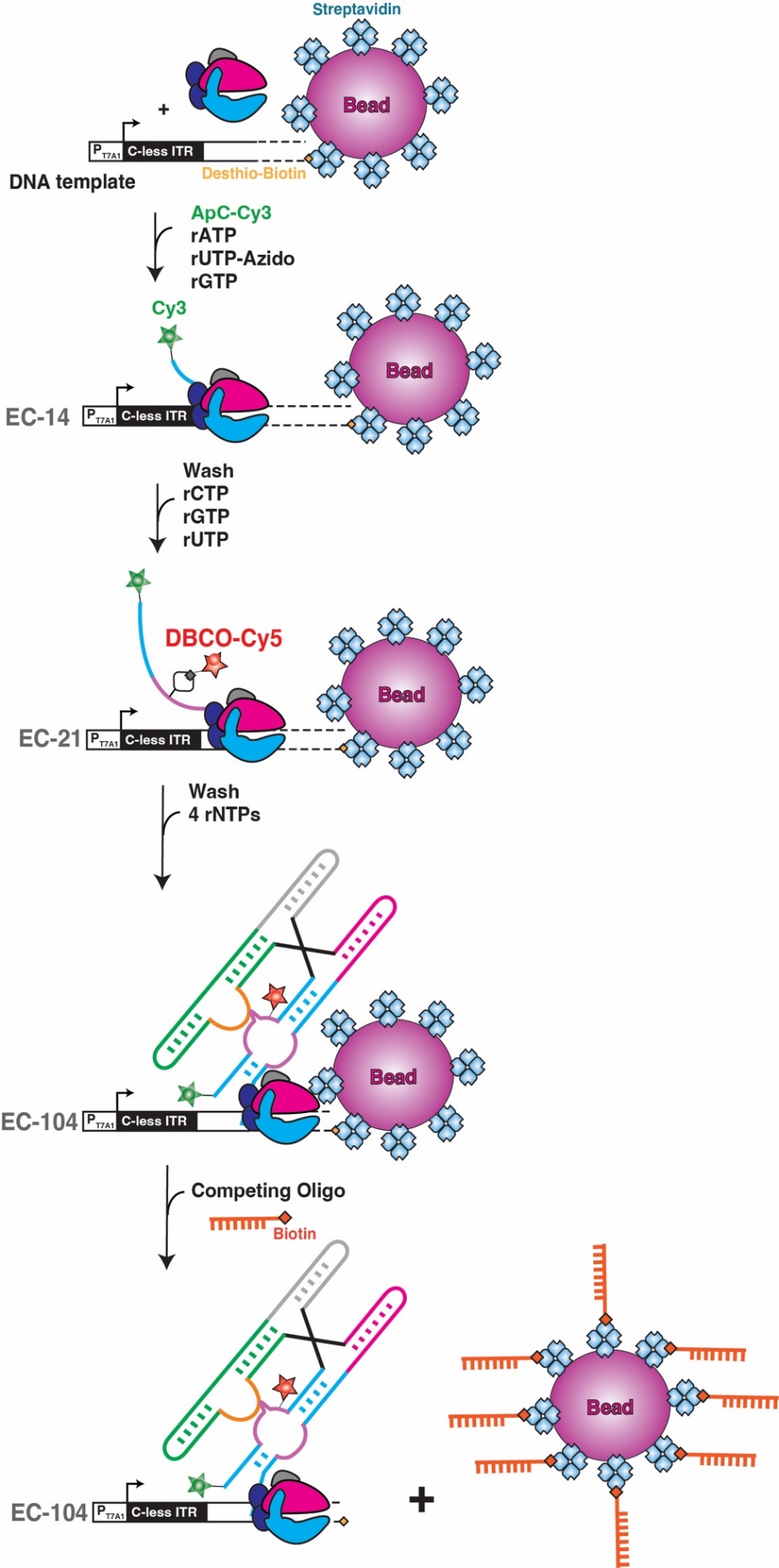

**Figure S12. Experimental procedure to generate a doubly labeled PEC-104**

The DNA template was immobilized on streptavidin-coated magnetic beads by labeling with a desthio-biotin moiety on the 3’end. Stepwise transcription was initiated with a 5’-Cy3-ApC dinucleotide and a subset of rNTPs lacking rCTP, leading to extension into EC-14. Incorporation of rUTP-azido is performed during this step labeling the one U in this sequence. After removing rNTPs from the complex, RNAP was walked through position 21 (EC-21) upon addition rGTP, rUTP and rCTP (lacking rATP). Labeling with DBCO-Cy5 was performed at this step. Next, the excess of unused fluorophores was washed away and all 4 rNTPs were added to reach the final roadblock position of PEC-104. Elution of the PEC from the beads was performed through incubation with a 20-fold excess of competing biotinylated oligonucleotide that replaces the weaker desthio-biotin ligand from streptavidin.

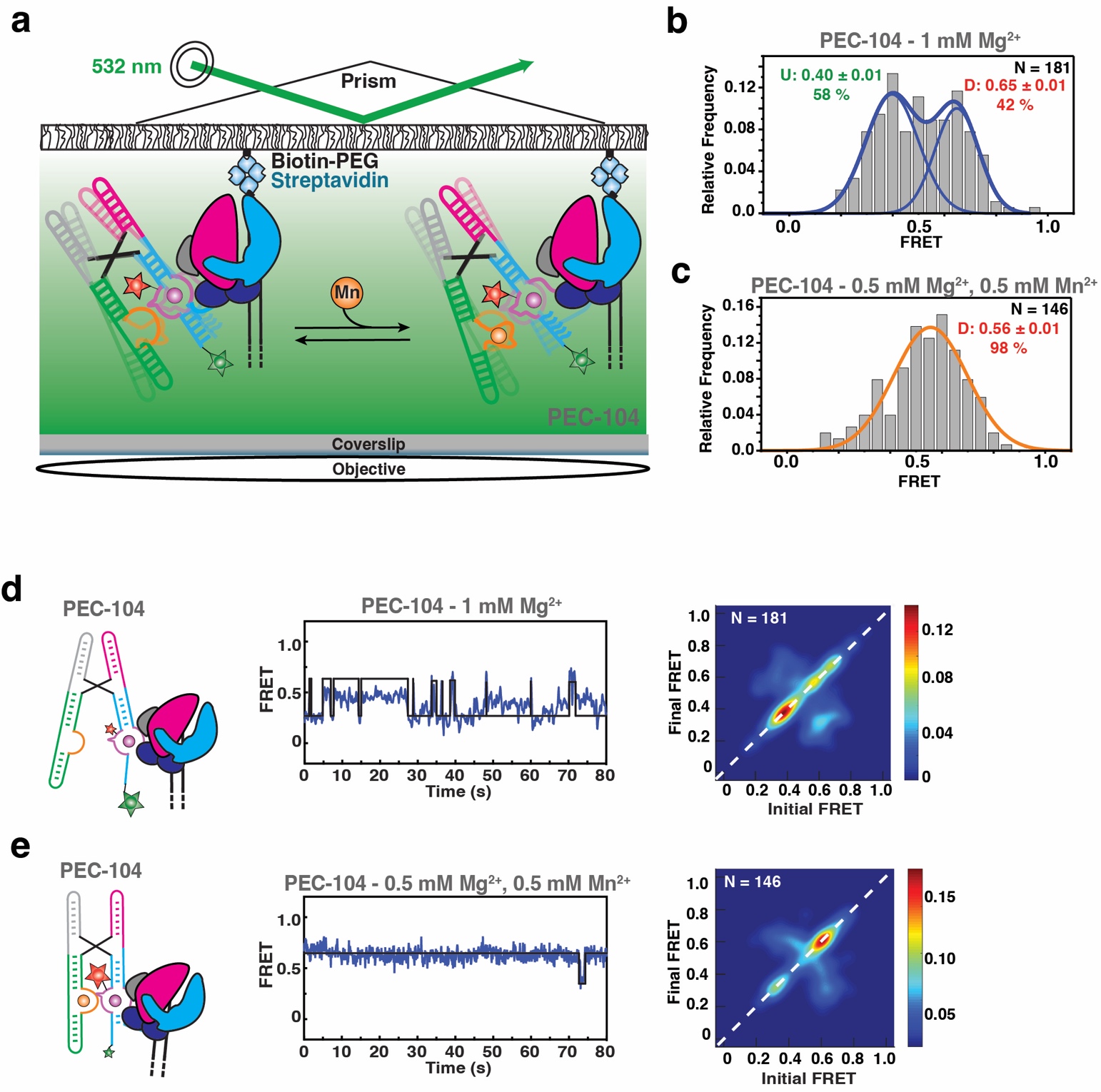

**Figure S13. Single molecule analysis of P1.1 folding in PEC-104**

(a) Experimental setup for smFRET interrogation of P1.1. PEC immobilization utilized the desthio-biotin moiety on the DNA that was also deployed to generate the transcription roadblock. Locations of donor (Cy3, green) and acceptor (Cy5, red) fluorophores are indicated.

(b, c) smFRET histograms of PEC-104 in the presence of only Mg^2+^ (b) and in the presence of both Mg^2+^ and Mn^2+^ (c). The mean FRET value with SD is reported, together with the total number of traces (N) included in each histogram.

(d, e) Representative FRET trajectories for PEC-104 in the presence of only Mg^2+^ (d) and in the presence of both Mg^2+^ and Mn^2+^ (e) along with the corresponding TODPs that represent dynamic traces as “off-diagonal” and static traces as “on-diagonal” contours, where the color scale shows the prevalence of each population.

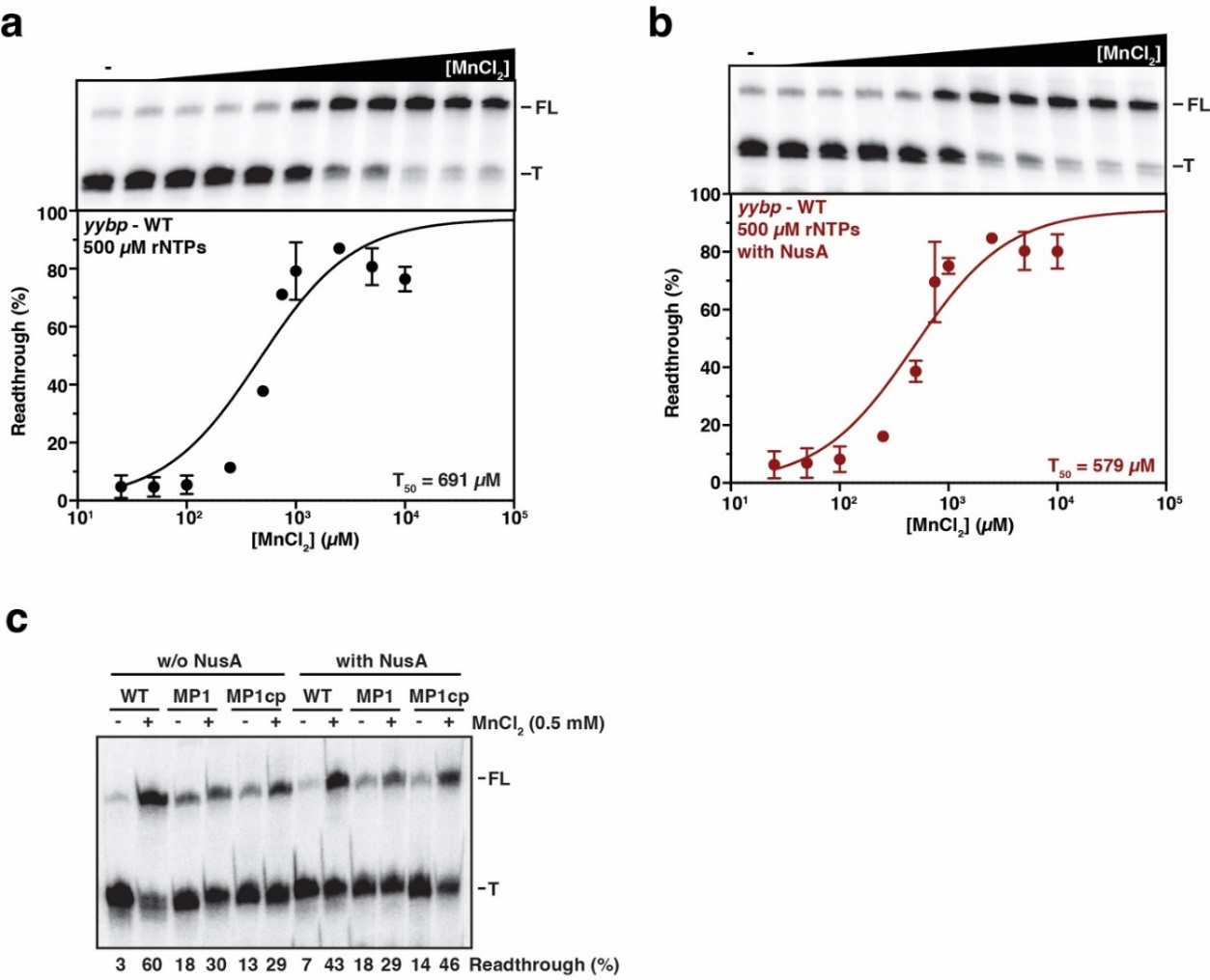

**Figure S14. Transcriptional regulation by the *yybp* riboswitch in the absence and presence of NusA**

(a, b) Mn^2+^-dependent single-round transcription assay of a DNA template containing the 5’-UTR of the *yybp* gene from *L. lactis* in the absence (a) and in the presence of 100 nM NusA transcription factor (b). Transcription reactions were performed using the *E. coli* RNAP with 500 µM rNTPs. The ^32^P labeled products were resolved on 6% polyacrylamide denaturing gel separating the full-length (FL) and terminated (T) RNA products. Plot of the fraction of transcription readthrough versus the concentration of MnCl_2_ is shown at the bottom. T_50_ values are indicated. Error bars represent the SD of the mean from independent replicates.

(c) Representative denaturing gel showing termination and antitermination in the absence (-) and presence (+) of Mn^2+^ in the context of the WT and mutants of the P1.1 helix. Transcriptions were performed in the absence or presence of 100 nM NusA factor with 25µM rNTPs. Percentages of readthrough are indicated at the bottom of the gel.

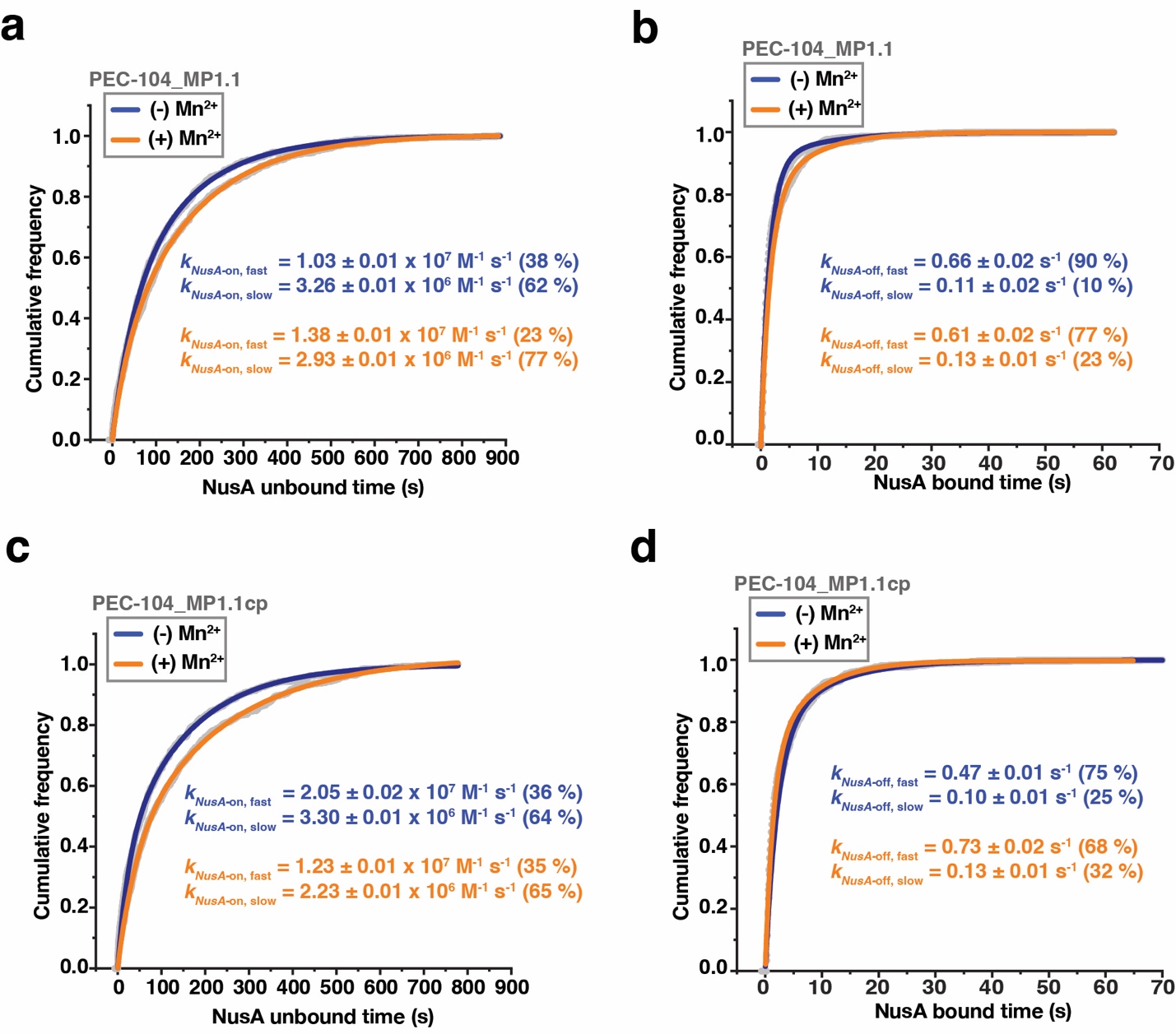

**Figure S15. Analysis of NusA binding to PEC-104**

(a, b) Plots displaying the cumulative unbound (a) and bound (b) dwell times of NusA-Cy5 to PEC-104 – MP1.1. Binding (*k*_on_) and dissociation (*k*_off_) rate constants of NusA-Cy5 are indicated.

(c, d) Plots displaying the cumulative unbound (c) and bound (d) dwell times of NusA-Cy5 to PEC-104 – MP1.1cp. Binding (*k*_on_) and dissociation (*k*_off_) rate constants of NusA-Cy5 are indicated. Total number of molecules analyzed for each condition is as follow: PEC-104 MP1.1 (-) Mn^2+^ = 232; PEC-104 MP1.1 (+) Mn^2+^ = 303; PEC-104 MP1.1cp (-) Mn^2+^ = 276; PEC-104 MP1.1cp (+) Mn^2+^ = 301.

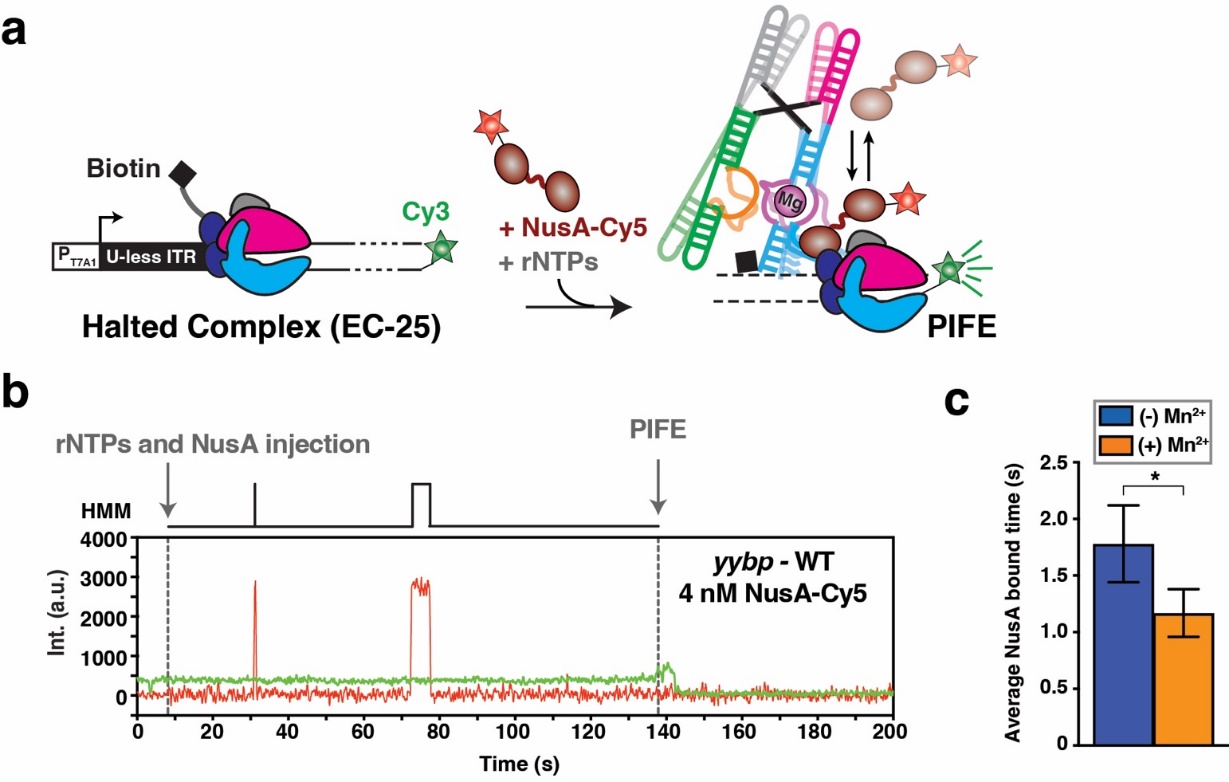

**Figure S16. Co-transcriptional ligand binding promotes NusA dissociation during transcription**

(a) Transcription of the DNA template labeled with Cy3 at the 3’-end using the *E. coli* RNAP allows the formation and the detection of a fluorescent halted complex (EC-12) attached to the microscope slide through the nascent RNA transcript. Transcription is restarted upon addition of all rNTPs, and addition of NusA-Cy5 in the transcription mix allows surveying co-transcriptional NusA binding in real time. When the RNAP reaches the end of the DNA template, the occurrence of the PIFE signal delimits the co-transcriptional window.

(b) Representative single-molecule trajectory showing the real-time transcription of the *yybp* riboswitch transcribed in the presence of 4 nM NusA-Cy5. Transcription restart is indicated by rNTP injection, and the end of transcription is indicated by PIFE. Repeated co-transcriptional bindings of NusA-Cy5 are monitored upon direct excitation of the Cy5 fluorescent dye. HMM is indicated on the top of the trace. (c) Overall bound time of NusA-Cy5 in the context of the *yybp* riboswitch transcription calculated in the absence (blue) and presence (orange) of 0.5 mM Mn^2+^ ion. Error bars are the SD from bootstrapping of all bound times collected. The total number of molecules analyzed in each dataset is as follows: (-) Mn^2+^ = 54; (+) Mn^2+^ = 45 (***p < 0.01, **p < 0.05, *p < 0.1).

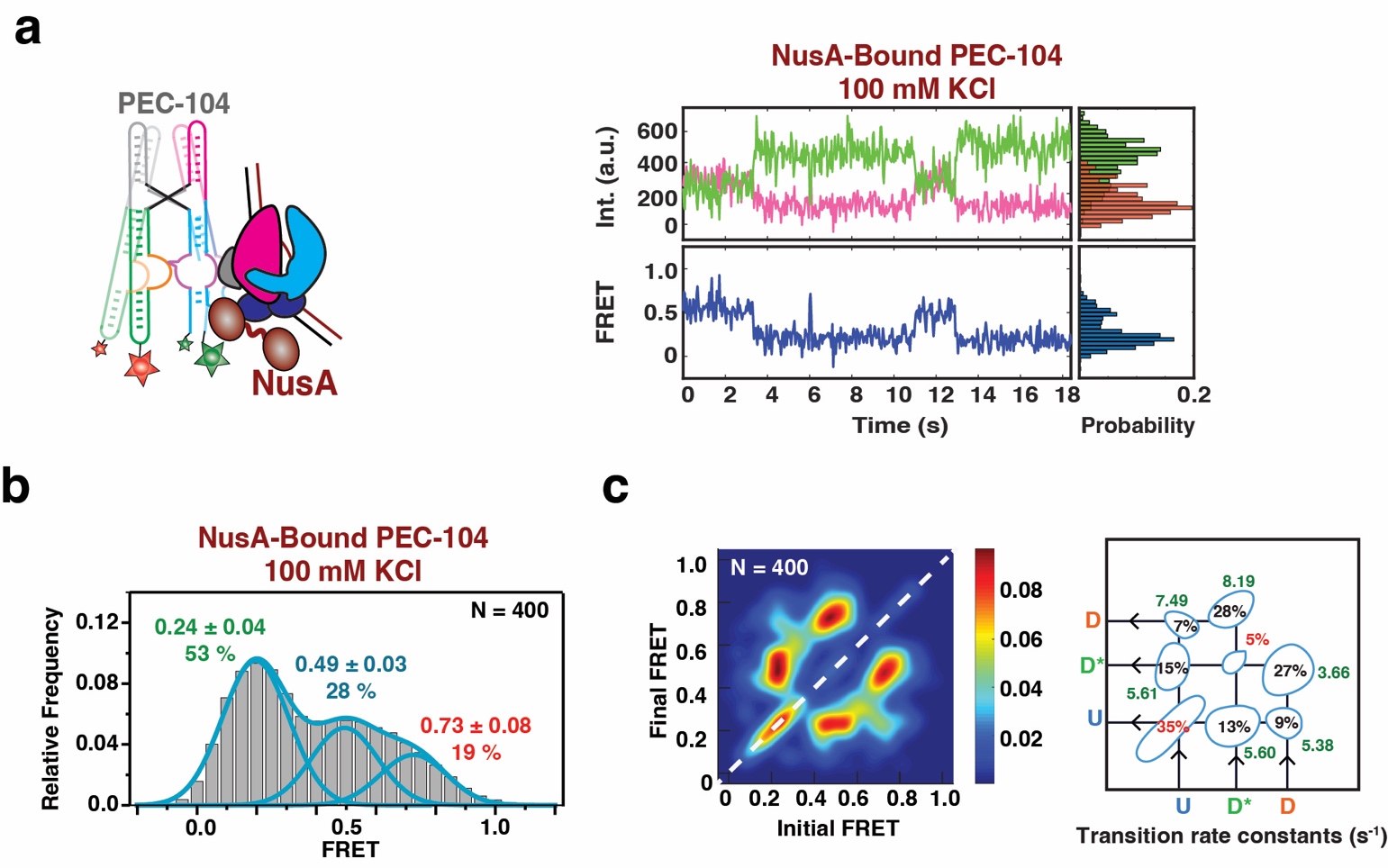

**Figure S17. smFRET analysis of the NusA-bound PEC-104 in the absence of divalent ion**

(a) Representative smFRET trace for NusA-bound PEC-104 in the absence of divalent ion (100 mM KCl). Green: Cy3, Pink: Cy5, Blue: FRET.

b) Population FRET histogram showing the equilibrium distribution of three FRET states under the condition in panel a.

(c) TODP showing the static and dynamic traces as “on-diagonal” and “off-diagonal” heat map contours, respectively for NusA-bound PEC-104. The color code indicates the fraction of each population. Transition rate constants for each transition between the different FRET states are indicated on the right.

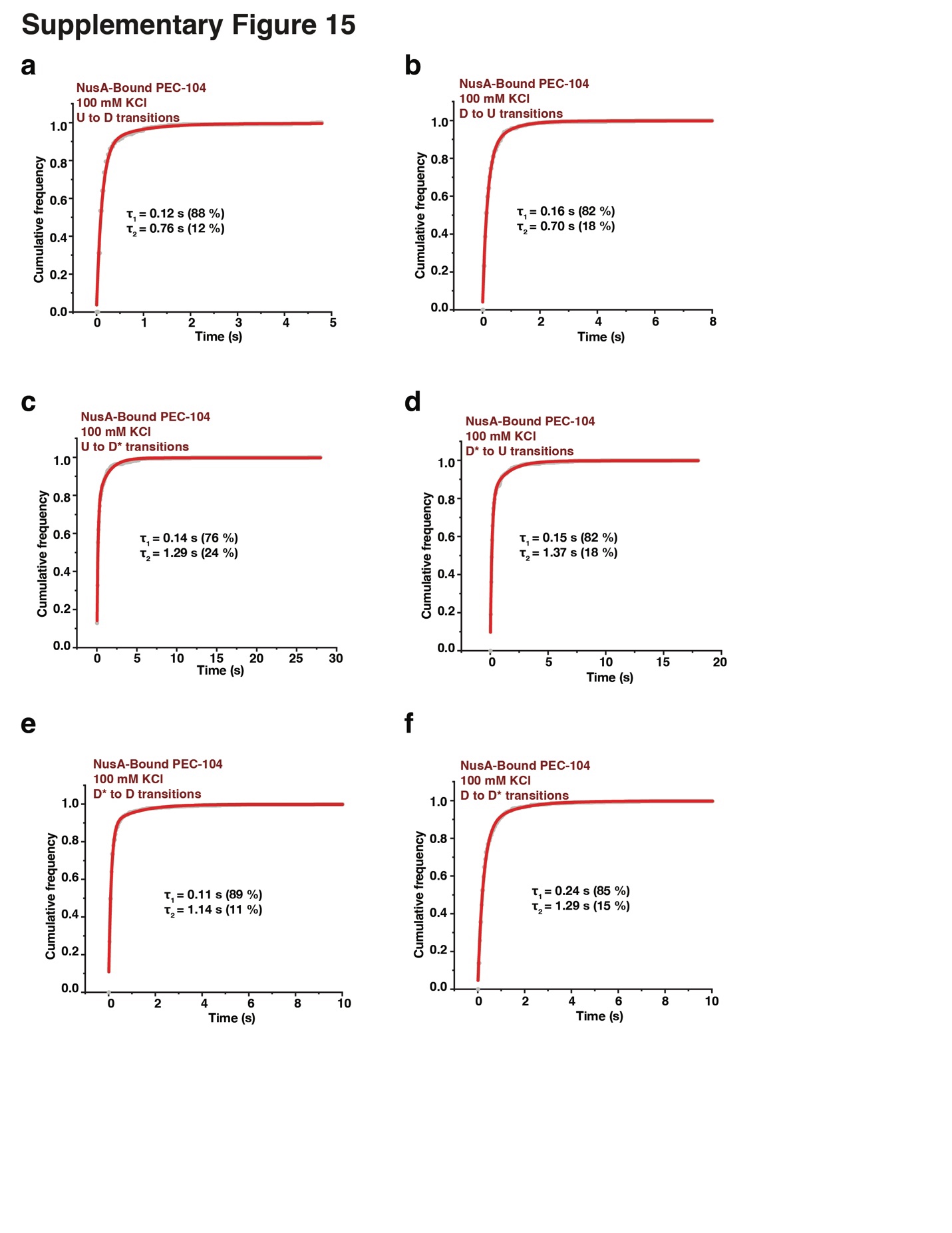

**Figure S18. Docking kinetics of NusA-bound PEC-104 in the absence of divalent ions**

(a-f) Cumulative dwell-time distributions of each transition between the three FRET states (U, D* and D) in the absence of divalent ions. The lifetimes and amplitudes of slow and fast components are also shown.

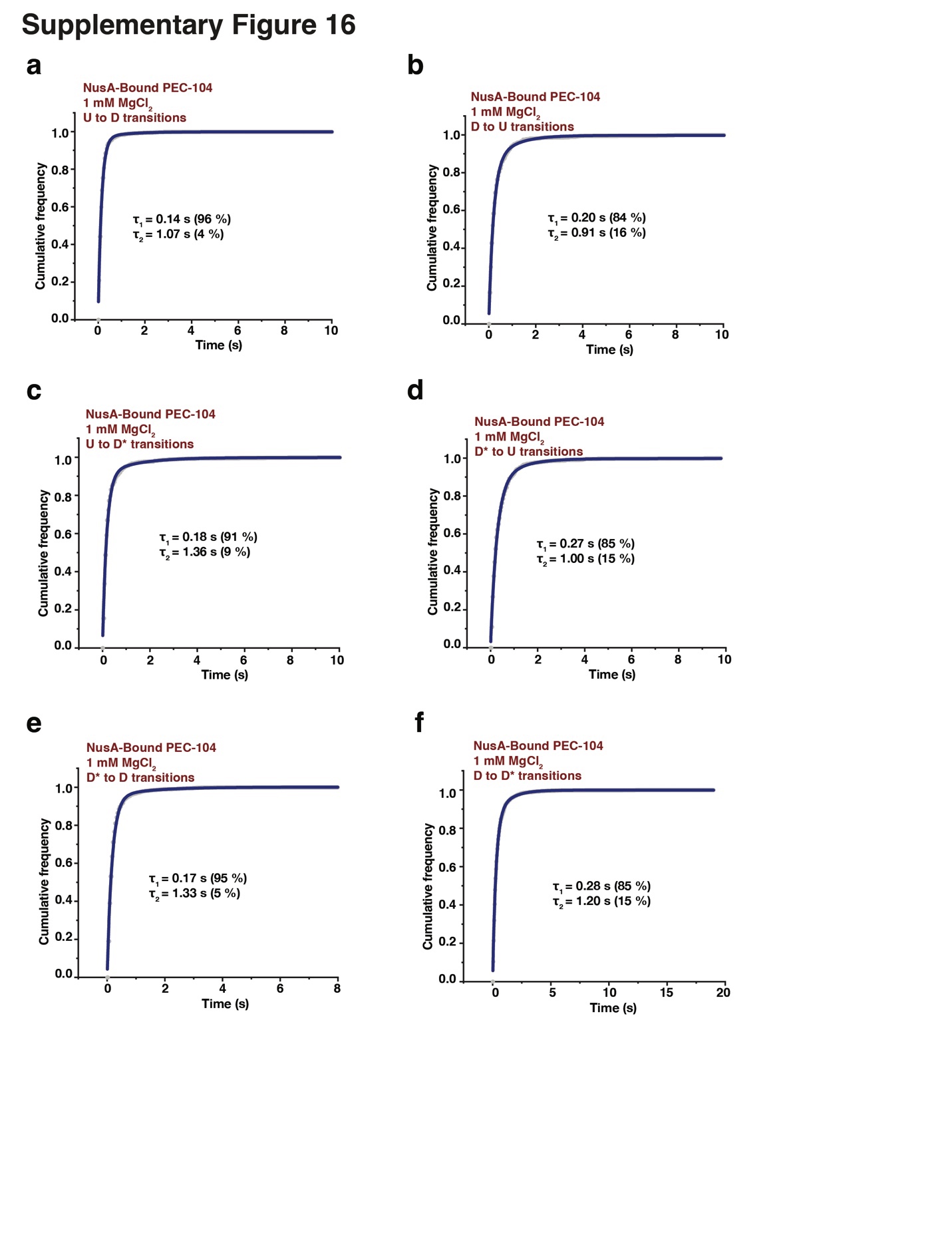

**Figure S19. Docking kinetics of NusA-bound PEC-104 in the presence of Mg^2+^ only**

(a-f) Cumulative dwell-time distributions of each transition between the three FRET states (U, D* and D) in the presence of 1 mM MgCl_2_. The lifetimes and amplitudes of slow and fast components are also shown.

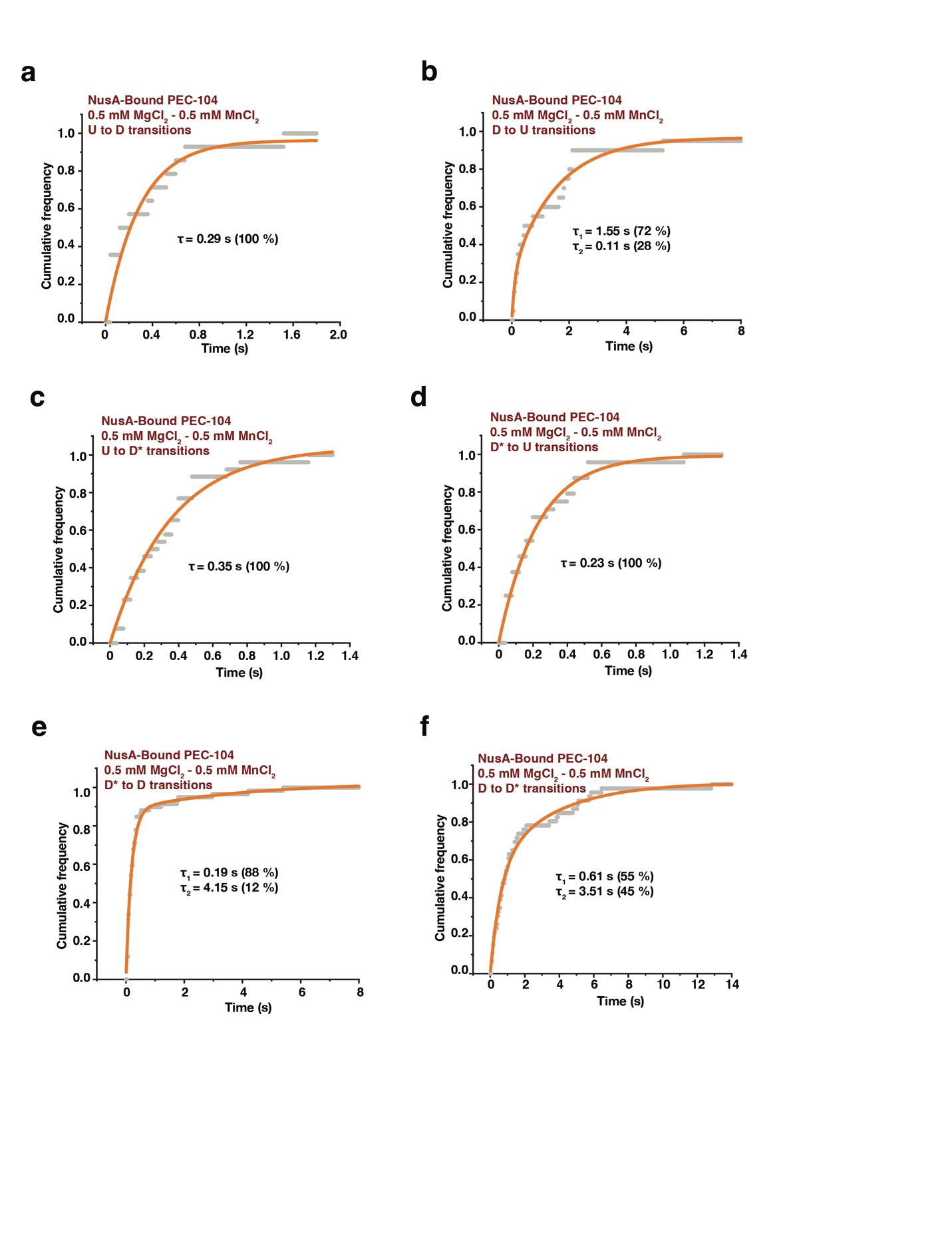

**Figure S20. Docking kinetics of NusA-bound PEC-104 in the presence of Mg^2+^ and Mn^2+^**

(a-f) Cumulative dwell-time distributions of each transition between the three FRET states (U, D* and D) in the presence of 0.5 mM MgCl_2_ and 0.5 mM MnCl_2_. The lifetimes and amplitudes of slow and fast components are also shown.

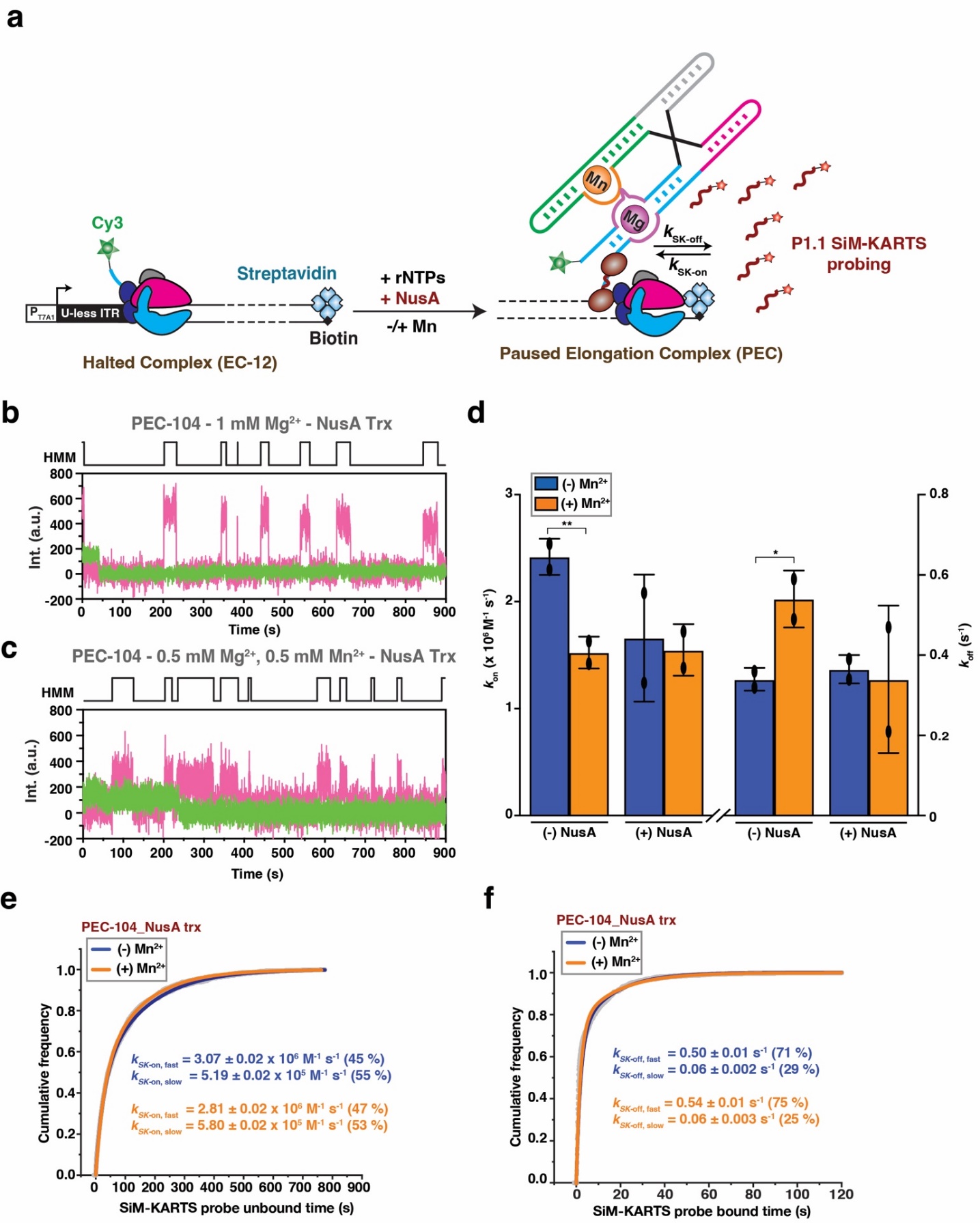

**Figure S21. SiM-KARTS analysis of PEC-104 in the presence of NusA during transcription**

(a) Fluorescently labeled PECs are transcribed in vitro using *E. coli* RNAP. The halted complex (EC-12) is prepared upon addition of a dinucleotide labeled with Cy3 (ApU-Cy3) and UTP deprivation (ATP/CTP/GTP) to halt the RNAP at the end of the U-less ITR. The biotin-streptavidin interaction at the 3’-end of the DNA template constitutes a stable transcriptional roadblock to stall the RNAP at the desired position.

(b, c) Representative single molecule trajectories showing the binding of the SiM-KARTS probe (red) to PEC-104 transcribed in the presence of NusA transcription factor with Mg^2+^ only (b) or with both Mg^2+^ and Mn^2+^ (c). HMM is indicated on the top of each trace.

(d) Overall binding rate constants (k_on_) of the SiM-KARTS probe in the contexts of PEC-104 transcribed in the absence (-) or in the presence (+) of 500 nM NusA transcription factor. Error bars are the standard deviation (SD) of the means from independent replicates.

(e, f) Plots displaying the cumulative unbound (e) and bound (f) dwell times of the SiM-KARTS probe to the P1.1 in the context of NusA-bound PEC-104. Binding (*k*_on_) and dissociation (*k*_off_) rate constants of the SiM-KARTS probe are indicated. Binding (*k*_on_) and dissociation (*k*_off_) rate constants of the SiM-KARTS probe are indicated. Total number of molecules analyzed for each condition is as follows: PEC-104 (-) NusA (-) Mn^2+^ = 350; PEC-104 (-) NusA (+) Mn^2+^ = 246; PEC-104 (+) NusA (-) Mn^2+^ = 319; PEC-104 (+) NusA (+) Mn^2+^ = 306.

**Table S1: Oligonucleotides used in this study.**

| **Oligonucleotide** | **Sequence (5’-3’)** |
| --- | --- |
| T7A1-PCR | TCCAGATCCCGAAAATTTATCAAAAAGAGTATTG |
| T7A1-Yybp-Start_AC | TCCAGATCCCGAAAATTTATCAAAAAGAGTATTGACTTAAAGTCTAACCTATAGGATACTTACAGCCACAAAGGGGAGTAGCGTCGG |
| Yybp-rev | CATAATGATTCTCCTTTATTTCTAACTGATATCATGAGAGG |
| Yybp-EC104 | /5Biosg/ATATCAAAACTGGCAAAGGTCTTGCTAAA |
| Yybp-EC104-DesthioBiotin | /5deSBioTEG/CT GGC AAA GGT CTT GCT AAA CAT ACA T |
| Yybp-probeSMK | /5Cy5/CCCTTTGT or 5Cy3/CCCTTTGT |
| Anchor_bio | /5Biosg/AGACCACGTTGAAAGATTGGGTTAC |
| Yybp-MP1.1 (2) | AAAACTGGCAAACCTCTTGCTAAACA |
| Yybp-MP1.1 (3) | TGTTTAGCAAGAGGTTTGCCAGTTTT |
| Yybp-G122C/G123C (2) | GGTTTTACAAAAAAAGAGGTCTGCCAGATATC |
| Yybp-G122C/G123C (3) | GATATCTGGCAGACCTCTTTTTTTGTAAAACC |
| tDNA-Yybp | CCAGTCATGCAGGACTGGCAAACCTTCTCTAAGTCTC |
| ntDNA-Yybp | GAGACTTAGAGAAGGAAATGGTTACCTGCATGACTGG |
| Yybp-EC104-MP1.1 | /5Biosg/ATATCAAAACTGGCAAACCTCTTGCTAAACATAC |
| Yybp-RT96 | AGACCACGTTGAAAGATTGGGTTACGGTCTTGCTAAACATACATAAGTATGTCAACAAAG |
| Yybp-MP1.1cp (2) | CCGACGCTACTCCGGTTTGTATGGCTGT |
| Yybp-MP1.1cp (3) | ACAGCCATACAAACCGGAGTAGCGTCGG |
| Yybp-RNA1 | /5Cy3/rUrCrArArArGrGrGrGrArGrUrArGrCrGrUrCrGrGrUrArArGrArCrCrGrArArArCrArArArGrUrCrGrUrCrArArUrUrCrGrUrGrArGrArU |
| Yybp-RNA2 | /5Cy5/rUrCrUrCrArCrCrGrGrCrUrUrUrGrUrUrGrArCrArUrArCrUrUrArUrGrUrArUrGrUrUrUrArGrCrArArGrArCrCrUrUrUrGrCrCrArG |
| 114_extension | /5Phos/rCrUrGrGrCrArGrArGrG |
| LNA-capture probe* | /5Biosg/TTTTTCC+TC+TGCC+AGC |

* LNA modifications are indicated with “+”

**Table S2: G104 pause half-lives measured in this study.**

| **Construct** | **Mn^2+^** | **τ (s) [pause half-life]*** |
| --- | --- | --- |
| Yybp-WT | - | 210 ± 5 |
|  | + | 775 ± 79 |
| Yybp-WT + NusA | - | 918 ± 35 |
|  | + | 983 ± 99 |
| Yybp-MP1.1 | - | 241 ± 1 |
|  | + | 227 ± 85 |
| Yybp-MP1.1 + NusA | - | 643 ± 70 |
|  | + | 665 ± 35 |
| Yybp-MP1.1cp | - | 137 ± 2 |
|  | + | 370 ± 38 |
| Yybp-MP1.1cp + NusA | - | 576 ± 54 |
|  | + | 646 ± 33 |

*The reported error is the standard deviation (SD) of the mean from independent replicates.

**Table S3: Kinetic parameters extracted from SiM-KARTS analysis.**

| **Construct** | **Mn^2+^** | ***k*_on_ (10^6^ M^-1^ s^-1^)** | ***k*_off_ (s^-1^)** |
| --- | --- | --- | --- |
| PEC-104 | - | Fast^a^: 3.67 ± 0.02 (59%)  Slow^a^: 0.63 ± 0.003 (41%)  Overall^b^: 2.42 ± 0.17 | Fast^a^: 0.55 ± 0.01 (61%)  Slow^a^: 0.08 ± 0.01 (39%)  Overall^b^: 0.34 ± 0.03 |
|  | + | Fast^a^: 2.93 ± 0.01 (44%)  Slow^a^: 0.49 ± 0.001 (56%)  Overall^b^: 1.53 ± 0.15 | Fast^a^: 0.36 ± 0.01 (85%)  Slow^a^: 0.06 ± 0.01 (15%)  Overall^b^: 0.54 ± 0.07 |
| PEC-104_MP1.1 | - | Fast^a^: 4.82 ± 0.05 (18%)  Slow^a^: 0.38 ± 0.001 (82%)  Overall^b^: 1.06 ± 0.04 | Fast^a^: 0.27 ± 0.01  Slow^a^: NA  Overall^b^: 0.30 ± 0.09 |
|  | + | Fast^a^: 3.52 ± 0.07 (8%)  Slow^a^: 0.45 ± 0.001 (92%)  Overall^b^: 0.68 ± 0.23 | Fast^a^: 0.34 ± 0.01  Slow^a^: NA  Overall^b^: 0.35 ± 0.04 |
| RT96 | - | Fast^a^: 4.55 ± 0.04 (51%)  Slow^a^: 0.29 ± 0.002 (49%)  Overall^b^: 2.44 ± 0.06 | Fast^a^: 0.46 ± 0.01 (86%)  Slow^a^: 0.04 ± 0.01 (14%)  Overall^b^: 0.46 ± 0.05 |
|  | + | Fast^a^: 5.35 ± 0.05 (50%)  Slow^a^: 0.30 ± 0.002 (50%)  Overall^b^: 2.45 ± 0.31 | Fast^a^: 0.44 ± 0.01 (88%)  Slow^a^: 0.04 ± 0.01 (12%)  Overall^b^: 0.37 ± 0.04 |
| PEC-104 + NusA | - | Fast^a^: 3.07 ± 0.02 (45%)  Slow^a^: 0.52 ± 0.002 (55%)  Overall^b^: 1.66 ± 0.59 | Fast^a^: 0.50 ± 0.01 (71%)  Slow^a^: 0.06 ± 0.002 (29%)  Overall^b^: 0.37 ± 0.04 |
|  | + | Fast^a^: 2.81 ± 0.02 (47%)  Slow^a^: 0.58 ± 0.002 (53%)  Overall^b^: 1.55 ± 0.24 | Fast^a^: 0.54 ± 0.01 (75%)  Slow^a^: 0.06 ± 0.003 (25%)  Overall^b^: 0.34 ± 0.18 |

^a^Values were calculated from single or double-exponential fits of the pool data from all the experiments in a given condition. The percentages indicate the contribution of each phase to the overall rate constant. The reported error is the standard deviation of the fit. In the case of single-exponential fit, only one value is reported arbitrarily as a fast rate constant.

^b^Values represent the average ± the standard deviation (SD) of the mean from independent replicates.

**Table S4: Percentage of stably docked (SD) traces observed in the smFRET analysis.**

| **Construct** | **static docked Traces** | **Total**  **traces** | **Percentage static docked (%)** |
| --- | --- | --- | --- |
| **PEC 104** |  |  |  |
| 100 mM KCl | 47 | 350 | 13 |
| 100 mM KCl with 1 mM Mg^2+^ | 63 | 319 | 19.7 |
| 100 mM KCl with 0.5 mM Mg^2+^ & 0.5 mM Mn^2+^ | 84 | 234 | 35.9 |
| **LNA 104** |  |  |  |
| 100 mM KCl | 14 | 122 | 11.5 |
| 100 mM KCl with 1 mM Mg^2+^ | 55 | 305 | 18 |
| 100 mM KCl with 0.5 mM Mg^2+^ & 0.5 mM Mn^2+^ | 149 | 326 | 45.7 |
| **LNA 114** |  |  |  |
| 100 mM KCl Only | 2 | 229 | 0.9 |
| 100 mM KCl with 1 mM Mg^2+^ | 20 | 258 | 7.8 |
| 100 mM KCl with 0.5 mM Mg^2+^ & 0.5 mM Mn^2+^ | 115 | 302 | 38.1 |

**Table S5: Kinetic parameters extracted from the NusA co-localization assay.**

| **Construct** | **Mn^2+^** | ***k*_on_ (10^6^ M^-1^ s^-1^)** | ***k*_off_ (s^-1^)** |
| --- | --- | --- | --- |
| PEC-104 (WT) | - | Fast^a^: 7.71 ± 0.01  Slow^a^: NA  Overall^b^: 7.05 ± 0.82 | Fast^a^: 0.93 ± 0.01 (93%)  Slow^a^: 0.13 ± 0.02 (7%)  Overall^b^: 0.83 ± 0.15 |
|  | + | Fast^a^: 3.97 ± 0.01  Slow^a^: NA  Overall^b^: 4.10 ± 0.11 | Fast^a^: 0.73 ± 0.01 (95%)  Slow^a^: 0.08 ± 0.01 (5%)  Overall^b^: 0.62 ± 0.18 |
| PEC-104 (MP1.1) | - | Fast^a^: 3.99 ± 0.01  Slow^a^: NA  Overall^b^: 3.97 ± 0.91 | Fast^a^: 1.03 ± 0.02 (93%)  Slow^a^: 0.17 ± 0.03 (7%)  Overall^b^: 0.89 ± 0.14 |
|  | + | Fast^a^: 4.10 ± 0.01  Slow^a^: NA  Overall^b^: 3.55 ± 0.93 | Fast^a^: 1.04 ± 0.02 (92%)  Slow^a^: 0.11 ± 0.03 (8%)  Overall^b^: 0.93 ± 0.04 |
| PEC-104 (MP1.1cp) | - | Fast^a^: 20.50 ± 0.10 (39%)  Slow^a^: 3.95 ± 0.01 (61%)  Overall^b^: 11.8 ± 1.76 | Fast^a^: 0.67 ± 0.01 (83%)  Slow^a^: 0.05 ± 0.01 (17%)  Overall^b^: 0.72 ± 0.17 |
|  | + | Fast^a^: 11.8 ± 0.05 (58%)  Slow^a^: 3.85 ± 0.02 (42%)  Overall^b^: 8.44 ± 1.05 | Fast^a^: 0.71 ± 0.01 (87%)  Slow^a^: 0.05 ± 0.01 (13%)  Overall^b^: 0.85 ± 0.08 |

^a^Values were calculated from single or double-exponential fits of the pool data from all the experiments in a given condition. The percentages indicate the contribution of each phase to the overall rate constant. The reported error is the standard deviation (SD) of the fit. In the case of single-exponential fit, only one value is reported arbitrarily as a fast rate constant.

^b^Values represent the average ± the standard deviation (SD) of the mean from independent experiments.
